# Contrasting megabenthic community structure across western Pacific seamounts to ∼5,000 m depth: Implications for conservation

**DOI:** 10.64898/2026.07.30.741913

**Authors:** Huang Jun-Long, Xun Lu, Dong Sun, Marie-Josee Fortin

## Abstract

**Aim:** Seamounts are key components of deep-sea ecosystems, yet how benthic community structure varies across depth gradients remains poorly understood. Here, we examine patterns of nestedness, turnover, and β-diversity within and among seamounts along the Kyushu–Palau Ridge (KPR) and assess their implications for conservation.

**Location:** Philippine Sea in the western Pacific.

**Major taxa studied:** Deep-sea megabenthic invertebrates (> 1 cm).

**Methods:** We surveyed six transects (13.3°N–22.9°N; 614–5,055 m depth; 165 km in total length) using a towed camera system, yielding 31,342 images and 14,218 identified individuals. Community structure was assessed through species depth-range overlap analysis to distinguish nestedness versus turnover patterns. β-diversity was partitioned into local contributions (LCBD) and species contributions (SCBD) to evaluate site-level uniqueness and species-level influence. Generalized linear mixed models were used to relate LCBD and SCBD to depth, slope, aspect, bathymetric position, and functional traits.

**Results:** Community structures varied among slopes and seamounts. Western, steeper slopes were dominated by nestedness, whereas gentler eastern slopes showed mixed patterns of nestedness and gradual turnover. Sessile species largely exhibited nestedness, while mobile taxa showed mixed structures. LCBD declined with depth overall but increased on north-facing slopes and sites with higher broad-scale bathymetric position. Wide-ranging generalist species contributed disproportionately to β-diversity.

**Main conclusions:** Megabenthic communities along the KPR showed contrasting patterns of nestedness and turnover across seamount slopes and depth gradients. Community variation was associated with depth and seafloor topography, while species with broader depth ranges contributed more strongly to β-diversity. The predominance of nestedness on western slopes suggests that shallow habitats may represent important components of seamount biodiversity, whereas the mixed patterns observed on eastern slopes highlight the contribution of both shallow and deeper, compositionally distinct habitats. These findings emphasise that conservation spatial planning for deep-sea seamount communities in tropical oligotrophic waters must account for both depth- and slope-specific variations, rather than relying on depth alone.

## Introduction

The deep ocean, once considered remote and largely inaccessible, is rapidly emerging as the next frontier for resource extraction (Levin *et al*., 2020). Driven by an escalating global demand for critical materials essential to energy industries, electronics, and defense, geopolitical interest has shifted toward the vast mineral deposits buried in the deep-sea ecosystems (Hein *et al*., 2013; Glover *et al*., 2026). This transition is not merely an economic endeavor but carries profound ecological consequences. Mining activities are poised to transform some of Earth’s least understood ecosystems at unprecedented scales, compounding long-standing pressures from climate change, pollution, and bottom-trawling (Ross *et al*., 2020; Jones *et al*., 2025; Victorero *et al*., 2025).

Among the most vulnerable ecosystems in the deep-sea landscape (> 200 m water depth) are seamounts (FAO, 2009; Schlacher *et al*., 2014; Watling & Auster, 2017; Swanborn *et al*., 2026). These towering undersea mountains rise hundreds to thousands of meters above the abyssal plains, creating rare islands of hard substrate in otherwise sediment-dominated environments (Rowden *et al*., 2010; Weber *et al*., 2025; Niyazi *et al*., 2026). Strong, persistent currents sweeping across their slopes support rich assemblages of suspension feeders, including corals and sponges, which in turn provide the structural complexity necessary for spawning grounds, nurseries, and refugia for numerous species (Genin *et al*., 1986; Clark *et al*., 2010; Shank, 2015). Such ecological features make seamounts vital to the structure and functioning of ocean ecosystems (Clark *et al*., 2010; Wang *et al*., 2024). Nevertheless, seamounts are also increasingly targeted for resource extraction because their slopes and surrounding areas can host polymetallic crusts and nodules (Du Preez *et al*., 2025). In this context, understanding biodiversity patterns on seamounts and their underlying drivers has become an urgent global priority (Clark *et al*., 2012).

Research across deep-sea ecosystems reveals pronounced gradients in species diversity and composition, shaped by a series of abiotic and biotic factors such as depth, latitude, energy, and competition (Rex *et al*., 1993; Levin *et al*., 2001; Woolley *et al*., 2016; Carter *et al*., 2025; O’Hara *et al*., 2025). Yet, seamount benthic communities defy simple generalizations. For example, early studies characterized seamounts as isolated “evolutionary islands” harboring high levels of endemism (Wilson Jr. & Kaufmann, 1987; de Forges *et al*., 2000), whereas later work found they often resemble adjacent continental slopes and may host fewer unique species than previously suggested (Samadi *et al*., 2006; O’Hara, 2007). Moreover, while bathymetric studies often identify a unimodal richness peak between 1,000 and 3,000 m (Brown & Thatje, 2014), recent observations in regions such as the Northeast Pacific, South Atlantic, and northern Gulf of Mexico have failed to conform to these established depth–diversity relationships (McClain *et al*., 2010; Bridges *et al*., 2022; Cordes *et al*., 2025). These inconsistencies likely reflect the limitations of prior research, which primarily relied on localized, small-scale surveys and focused solely on a few indicator taxa (Cho & Shank, 2010; McClain *et al*., 2010; Shen *et al*., 2021; Watling & Auster, 2021; Lu *et al*., 2024). More importantly, these knowledge gaps create a fundamental dilemma for conservation and management (Clark *et al*., 2012): Should conservation measures target local endemics or adopt a regional approach if seamounts function as dispersal stepping stones across ocean basins? Should efforts prioritize species-rich “source” habitats or design conservation networks spanning multiple depth zones to capture distinct, stratified assemblages, and which areas warrant conservation priority?

Addressing these questions requires moving beyond α-diversity (i.e., within-site diversity) to examine the deeper fabric of community structure at large spatial scales, particularly patterns of species’ depth-range overlap and β-diversity (i.e., the variation in community composition across sites). These measures allow us to distinguish between two contrasting patterns of community structure: nestedness or turnover (Cartes & Carrassón, 2004; McClain & Rex, 2015). High nestedness suggests that assemblages in species-poor sites are just a subset of those in species-rich sites, such that protecting a few species-rich sites could capture most biodiversity. In contrast, turnover-dominated pattern indicates species replacement along depth gradients, necessitating conservation networks that encompass multiple depth zones to preserve distinct communities (Baselga, 2010). Furthermore, established β-diversity frameworks also enable partitioning of variance in community data by the contributions of individual species and sites (Legendre & De Cáceres, 2013). Disentangling these patterns and pinpointing biological assets that require priority protection can therefore provide critical insights for designing effective conservation strategies for vulnerable seamount ecosystems.

The Kyushu–Palau Ridge (KPR) is the longest seamount chain in the western Pacific, a region that hosts the highest concentration of deep-sea seamounts globally (Yesson *et al*., 2011). Large portions of the KPR have been designated as Ecologically or Biologically Significant Marine Areas under the Convention of Biological Diversity (CBD, 2016). Despite this recognition, the region’s deep-sea biota still remains understudied (Lu *et al*., 2024). To help inform conservation planning, the DY59 cruise aboard the research vessel *Dayanghao*, organized by the China Deep Ocean Affairs Administration, conducted a two-month survey between July and August 2020 to collect biological and environmental data from seamounts along the KPR. We documented deep-sea benthic communities through six transects (VL01–VL06) across four seamounts spanning 13.3°N to 22.9°N (Figure 1). Using a towed camera system (Figure S1), we surveyed depths from 614 m to 5,055 m (Figure S2A). Based on these data, we characterized the spatial ecology of megabenthic communities along the KPR depth gradient with two complementary approaches: species depth-range overlap analysis and the β-diversity partitioning framework (Pielou, 1977; Legendre & De Cáceres, 2013), and discussed the implications of the observed pattern for conservation planning. Specifically, we asked: (ⅰ) Does community structure follow a nestedness- or turnover-dominated pattern, and are the inferred patterns consistent among transects located on different seamounts and on contrasting slope aspects, or between sessile and mobile taxa? (ⅱ) Which sites contribute most to β-diversity within each transect, and how is the magnitude of these local contributions related to depth and other topographic factors? and (ⅲ) Which species most strongly drive variation in community composition across depth zones, and can their depth ranges and/or functional traits (e.g., sessility, size, feeding habit) predict their contributions?

**Figure 1.**
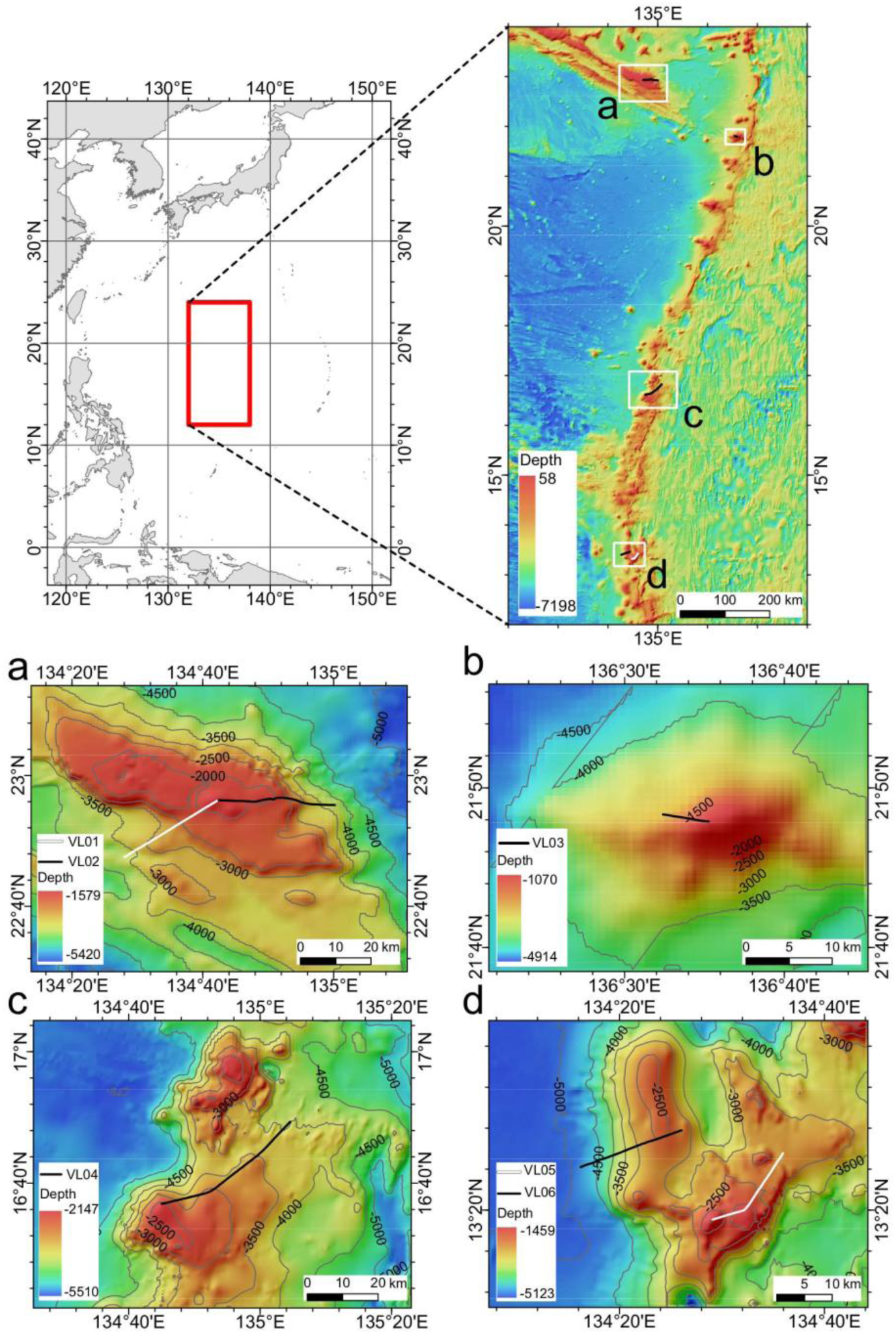
Study area and locations of six seamount survey transects VL01–VL06. Panel (a) VL01 & VL02: the eastern segment of the Oki-Daito Ridge; panel (b) VL03: Kasei Bank; panel (c) VL04: Mugiboshi Seamount; panel (d) VL05 & VL06: Chetiruir Seamount.

## Materials and Methods

### Study area

The Kyushu–Palau Ridge (KPR) is the longest seamount chain in the western Pacific Ocean, extending north–south across the Philippine Sea. This ridge exhibits highly complex and rugged topography, with discontinuous seamounts separated by latitudinal gaps near 18°N, 23°N, 24°N, and 25°N (Qin *et al*., 2021). The KPR is typically divided into three segments: northern (23°N to Kyushu Island), central (15°N–23°N), and southern (south of 15°N). Large portions of the northern and central KPR have been designated as Ecologically or Biologically Significant Marine Areas (EBSAs) under the Convention on Biological Diversity due to their exceptional biodiversity, ecological productivity, and role as spawning grounds for species such as sea eels (CBD, 2016). Despite this recognition, the deep-sea fauna of the region remains poorly studied (Lu *et al*., 2024). Here, we focused on four seamounts located in the central–southern KPR, including three deep-water seamounts (summit depth >800 m) and one intermediate seamount (summit depth 200–800 m), spanning latitudes from approximately 13.3°N to 22.9°N.

### Biological data collection

Biological data collection was conducted during the DY59 cruise aboard the research vessel *Dayanghao* between July and August 2020. Sampling was conducted on six transects (VL01–VL06) across four seamounts, covering a total of 165 km of deep-sea camera survey transects at depths from 614 m to 5,055 m. Transects were positioned to capture variability across seamount slopes: VL01 and VL02 extended westward and eastward on the same seamount, VL05 (eastern slope) and VL06 (western slope) formed a similar pair on another, while VL03 and VL04 were located on the western and eastern slopes of two different seamounts, respectively (see Table 1 for detailed transect profiles).

**Table 1.** Geographic and sampling characteristics of the survey transects. Transects on the same seamount are labeled with the same color.

| Transect | Seamount | Slope | Longitude (° E) | Latitude (° N) | Depth range | # Plots | # Observed individuals |
|---|---|---|---|---|---|---|---|
| VL01 | Oki-Daito Ridge | Western | 134.47–134.70 | 22.79–22.94 | 1493–3877 | 59 | 1207 |
| VL02 | Oki-Daito Ridge | Eastern | 134.71–135.00 | 22.92–22.94 | 1506–3693 | 62 | 2616 |
| VL03 | Kasei Bank | Western | 136.54–136.59 | 21.80–21.81 | 614–2253 | 11 | 1262 |
| VL04 | Mugiboshi Seamount | Eastern | 134.75–135.08 | 16.61–16.82 | 2061–4300 | 87 | 5895 |
| VL05 | Chetiruir Seamount | Eastern | 134.49–134.60 | 13.32–13.43 | 1039–2913 | 38 | 2599 |
| VL06 | Chetiruir Seamount | Western | 134.27–134.43 | 13.40–13.46 | 1998–5055 | 41 | 639 |

We mapped all transects onto the General Bathymetric Chart of the Oceans (GEBCO) gridded bathymetry data, which provides a high-reliability global bathymetric dataset at a spatial resolution of 15 arc-seconds (GEBCO Bathymetric Compilation Group, 2025). Considering the orientation of the transects, the mean transect length corresponding to a single GEBCO grid cell in our study area was approximately 500 m. For consistency with the spatial resolution of the GEBCO bathymetry data and to facilitate subsequent data processing, we divided all transects into 500 m segments, whose midpoints were then used as the location of the corresponding sampling plot. We further checked the spatial alignment between sampling plots and the gridded bathymetry data and found that each sampling plot was well matched to a distinct GEBCO grid cell.

A towed camera system equipped with high-definition video, digital still cameras (image resolution: 2560 × 1920 pixels), illumination lamps, an altimeter, and dual laser beams was deployed to capture images of benthic habitats (Figure S1). During the surveys, the system was towed at a constant speed of 1.0 knot (1.852 km/h). To ensure accurate real-time positioning, it carried an underwater transponder and was tracked using a shipborne POSIDONIA II ultra-short baseline (USBL) system (iXBLUE Inc., France). During each survey, the camera platform was maintained at an altitude of approximately 3 m above the seafloor. At this height, the digital still camera automatically captured images at 6-second intervals, yielding an average seabed coverage of 6 m² per photograph. In parallel, continuous 4K high-definition video footage was recorded throughout the transects. Following the predefined plot-partitioning procedure, we originally obtained 330 plots. After quality control, we retained 298 valid plots and excluded the remaining 32 plots for subsequent analyses, mainly due to poor positioning data quality, low-resolution imagery, and unstable camera altitude above the seafloor caused by complex terrain.

For post-survey analyses, a team of taxonomists examined all still photographs and identified megabenthic organisms to the lowest possible taxonomic level (or to distinct morphological species; Figure 2). The average (*±* SD) number of still images analyzed per valid plot (*n* = 298) was 105 ± 64. All quantitative biological data used in this study were derived exclusively from counts of megabenthic individuals recorded in these images. The continuous 4K video footage was not used for quantitative enumeration; instead, it served as an auxiliary resource that provided dynamic views and additional morphological details to support accurate taxonomic identification. Taxonomic assignments followed the World Register of Marine Species (WoRMS) guidelines (WoRMS Editorial Board, 2026). Organisms that could not be reliably identified were excluded from quantitative analyses. Fishes and crustaceans were also omitted due to their high mobility, as they either rapidly moved away from or persistently followed the towed camera system.

**Figure 2.**
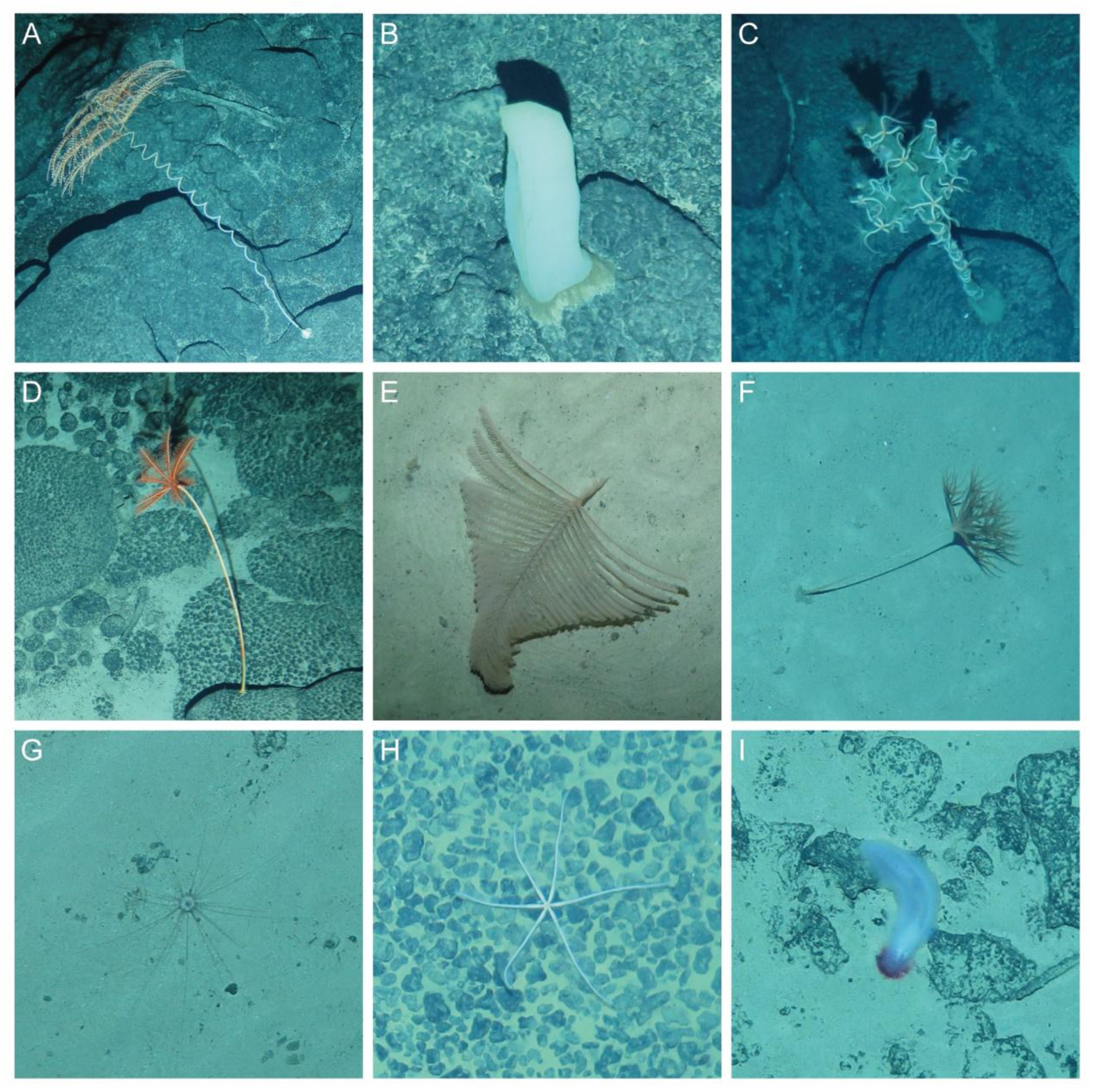
Examples of collected images. A: *Iridogorgia* spp. indet. (observed in VL01, 1720 m); B: *Poliopogon* sp.1 indet. (VL04, 3538 m); C: *Ophioplinthaca* spp. indet. (VL01, 1600 m); D: *Proisocrinus ruberrimus* (VL01, 1730 m); E: *Schizopathes* spp. indet. (VL03, 2083 m); F: *Umbellula* spp. indet. (VL03, 1756 m); G: *Plesiodiadema globulosum* sp. inc. (VL03, 1730 m); H: *Freyastera* spp. indet. (VL04, 3847 m); I: *Benthodytes sanguinolenta* sp. inc. (VL05, 1877 m). VL01–VL05 refer to the survey transect, see details in Table 1. Photo credit: Dong Sun.

Biological traits, including body size, mobility, feeding mode, and growth form, were compiled from the Marine Life Information (MarLIN, 2006) and WoRMS online databases (WoRMS Editorial Board, 2026), published literature, and direct observations (Table 2). These traits were selected because of their strong associations with deep-sea ecosystem functioning and species’ sensitivity to physical disturbances (Beauchard *et al*., 2017; González-Irusta *et al*., 2018; de la Torriente *et al*., 2020; Shen *et al*., 2024). When trait data were unavailable, information was inferred from closely related taxa.

**Table 2.** Biological traits used in this study.

| Traits | Category | Description |
|---|---|---|
| Movement | Sessile | Attach to hard or soft seafloor or other organisms, unable to move |
|  | Mobile | Non-sessile, able to move |
| Body size | ≤ 5 cm | Body length ≤ 5 cm |
|  | 5–50 cm | Body length > 5 cm and ≤ 50 cm |
|  | > 50 cm | Body length > 5 cm |
| Living habit | Solitary | Able to live alone |
|  | Associated | Prefer to attach to other organisms |
| Feeding habit | Suspension feeder | Passively trap waterborne food particles, including live prey, using external body structures |
|  | Filter feeder | Passively trap waterborne food particles, including live prey, using external body structures |
|  | Deposit feeder | Ingest sediments and food particles on or in the seabed |
|  | Predator | Actively prey on live animals or consume dead animals |
|  | Grazer | Remove attached organic particles from hard or soft substrata using abrasive mouthparts |
| Body form | Flattened | Flattened body form |
|  | Globular or mound | Globular or mound body form |
|  | Stalk | Erect or stalk body form |
|  | Topological | Topological or tree-like body form |

### Environmental data

As the KPR is located in an oligotrophic region and most sampling plots were below the permanent thermocline, we expected their environmental chemical properties to be highly stable and homogeneous (Sanders & Hessler, 1969; Wang *et al*., 2024). Consequently, we focused our analyses on topographic factors, which represent the primary source of environmental variation in this region. The GEBCO gridded bathymetric data were processed using the Benthic Terrain Modeler in ArcGIS to derive seabed characteristics (Walbridge *et al*., 2018). Depth measurements from the towed camera’s pressure sensors were integrated with vessel GPS data to georeference each transect’s site. The final environmental dataset included longitude, latitude, depth, slope, curvature, roughness, and broad-scale and fine-scale bathymetric position indices (BBPI and FBPI). Curvature, defined as the second spatial derivative of seabed topography, quantifies the degree of seafloor undulation (Yan *et al*., 2024). Roughness, calculated as the ratio of actual surface area to its planar projection, characterizes terrain complexity (Lundblad *et al*., 2006). Bathymetric Position Indices describe the relative position of each grid cell within the surrounding landscape, in which positive values indicate relatively elevated areas, such as bumps or ridges, whereas negative values indicate relatively depressed areas, such as depressions or valleys (Lundblad *et al*., 2006). Following Lundblad *et al*. (2006) and Fan *et al*. (2022), FBPI grids were generated using an outer radius of 2 grid cells and inner radius of 1 grid cell, whereas BBPI grids used an outer radius of 4 grid cells and inner radius of 1 grid cell.

### Species depth-range overlap analysis

We analyzed species range overlap along depth gradients to determine whether community structure was dominated by turnover or nestedness. For each transect, species ranges were defined by their first and last occurrences on the depth gradient. All possible species pairs were classified into three categories: non-overlapping, partially overlapping, or completely overlapping. These counts were summarized in the overlap triplet *L* = (*ω_none_*, *ω_partial_*, *ω_complete_*), where *ω_none_*, *ω_partial_*, and *ω_complete_* represent the number of non-overlapping, partially overlapping, and completely overlapping pairs, respectively (Pielou, 1977). This approach focuses on presence–absence data and captures structural signals without requiring abundance information.

To evaluate whether the observed overlap pattern differs from random expectation, we used the random-order null model (*H*_0_), which assumes that all species boundaries occur in a fully random sequence along the gradient, subject to the constraint that each species’ lower boundary follows its own upper boundary (Pielou, 1977). For *n* species, the expected overlap triplet under *H*_0_ is:

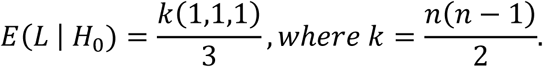

We computed deviations ***L*** – E(***L*** | *H*_0_) for each transect, and their signs provide clues to infer community structures. For example, a sign pattern of (−, −, +) indicates an overabundance of completely overlapping pairs that implies a nestedness-dominated community structure, whereas a (+, −, −) pattern indicates an excess of non-overlapping pairs, suggesting a turnover-dominated structure (Fortin & Dale, 2014).

### β-diversity partitioning

To assess how individual sites and species contribute to overall β-diversity along depth gradients, we applied the variance-partitioning framework proposed by Legendre and De Cáceres (Legendre & De Cáceres, 2013). In this approach, β-diversity is expressed as the total variance in community composition across sites. We first applied a Hellinger transformation to the species abundance matrix to reduce the influence of double zeros and make Euclidean distances ecologically meaningful for community data (Legendre & Gallagher, 2001). From the transformed matrix, we computed two complementary metrics: Local Contributions to Beta Diversity (LCBD) and Species Contributions to Beta Diversity (SCBD). LCBD quantifies how compositionally unique each site is compared to all others; sites with high LCBD values represent areas that disproportionately shape overall β-diversity. SCBD measures the extent to which each species varies in occurrence or abundance across sites, highlighting species that drive compositional heterogeneity (Legendre & De Cáceres, 2013; Lu *et al*., 2024; Tapia-Guerra *et al*., 2026). Both metrics are derived from the decomposition of total variance: LCBD values correspond to the relative share of each site in the total sum of squares, while SCBD values reflect the relative share of each species. These metrics allow us to identify ecologically distinctive sites and the influential species within the depth gradient. Calculations were performed using the *adespatial* packages (Dray *et al*., 2025).

### Statistical analysis

All statistical analyses were conducted in R version 4.5.0 (R Core Team, 2025), and visualizations were created using the *ggplot2* package (Wickham, 2016).

To examine how LCBD varied with depth, we performed Pearson correlation analyses both within each transect and across all transects combined. To further evaluate the effects of topographic factors on LCBD, we fitted a generalized linear mixed model with a beta distribution, as LCBD values are bounded by [0, 1], and included transect as a random factor. The aspect variable was transformed using a sine function to represent north–south orientation, and all other predictors were standardized to a mean of 0 and a standard deviation of 1 prior to analysis.

To assess whether generalists, specialists, or rare species contributed most strongly to β-diversity, we calculated Pearson correlations between SCBD and species’ depth-range extent for each transect. For species occurring in multiple transects with varying SCBD values, we used the average SCBD. Lastly, to investigate how life-history and morphological traits influenced SCBD, we constructed a generalized linear model relating SCBD to species traits. All trait variables were categorical and were therefore encoded as dummy variables in the model. Mixed-effect models were built and analyzed using the *glmmTMB* packages (Brooks *et al*., 2017).

## Results

### Species richness and community composition across transects

The sampling efforts yielded 31,342 high-resolution images and 14,218 megabenthic individuals identified from these images across the 298 valid plots. Overall, we recorded 122 (morpho)species representing 91 genera, 67 families, 32 orders, and 8 classes. At the class level, Anthozoa accounted for the highest proportion of individuals in transects VL02 (40.7%), VL03 (51.8%), VL04 (43.5%), and VL06 (26.1%), followed by Ophiuroidea. Conversely, Ophiuroidea dominated VL01 (40.8%) followed by Anthozoa, while Crinoidea was most abundant in VL05 (44.6%).

### Contrasting community structures within and between seamounts

Transects VL01 and VL02 exhibited markedly different patterns of community structure despite being located on the same seamount. VL01 had an observed overlap triplet of (442, 463, 1111), which, when compared with the expected values of (672, 672, 672), yielded a (–, –, +) signature. This pattern reflects an overrepresentation of completely overlapping pairs, indicating a strongly nested community structure. VL02, however, exhibited an observed triplet of (119, 884, 1412) relative to expected values of (805, 805, 805), yielding a (–, +, +) pattern. This represents an excess of both partially and completely overlapping pairs, suggesting a mixed structure of both gradual turnover and nestedness (Figure 3). In contrast, transects VL05 and VL06 on a different seamount both exhibited nestedness-dominated structures, as did VL03. Lastly, VL04 resembled VL02, displaying a mixture of gradual turnover and nestedness (Table S1).

**Figure 3.**
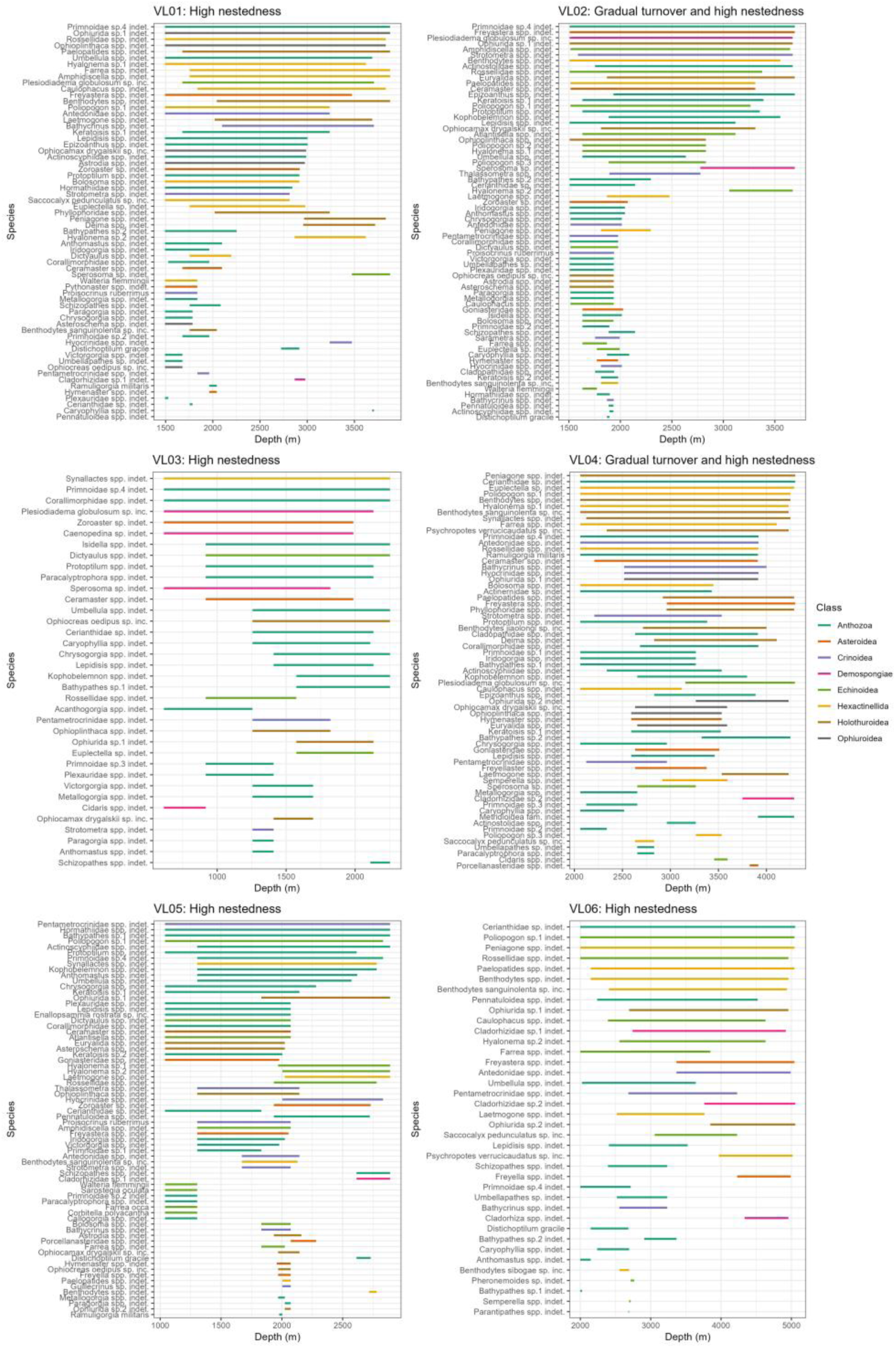
Depth-range patterns of species, arranged by the depth-range extent and maximum depth of occurrence within each transect (VL01–VL06; see Table 1). Species from different taxonomic classes are shown in distinct colors.

We further performed depth-range overlap analyses separately for sessile and mobile species per transect (Table S1). Patterns of sessile species generally mirrored those observed for the full community and exhibited highly nested patterns, except in VL02. However, mobile species predominantly showed mixed structures combining gradual turnover with nestedness in four of the six transects.

### Relationships between LCBD and environmental variables

Across sampling sites, LCBD values ranged from 0.009 to 0.131. LCBD was positively correlated with depth in VL02 (*R* = 0.70, *p* < 0.001) and VL04 (*R* = 0.28, *p* = 0.01), and negatively correlated with depth in VL03 (*R* = –0.64, *p* = 0.033) and VL06 (*R* = –0.73, *p* < 0.001). No significant relationships were detected for VL01 and VL05 (Figure 4A). Across all transects combined, LCBD was negatively correlated with depth (*R* = –0.42, *p* < 0.001; Figure 4B).

**Figure 4.**
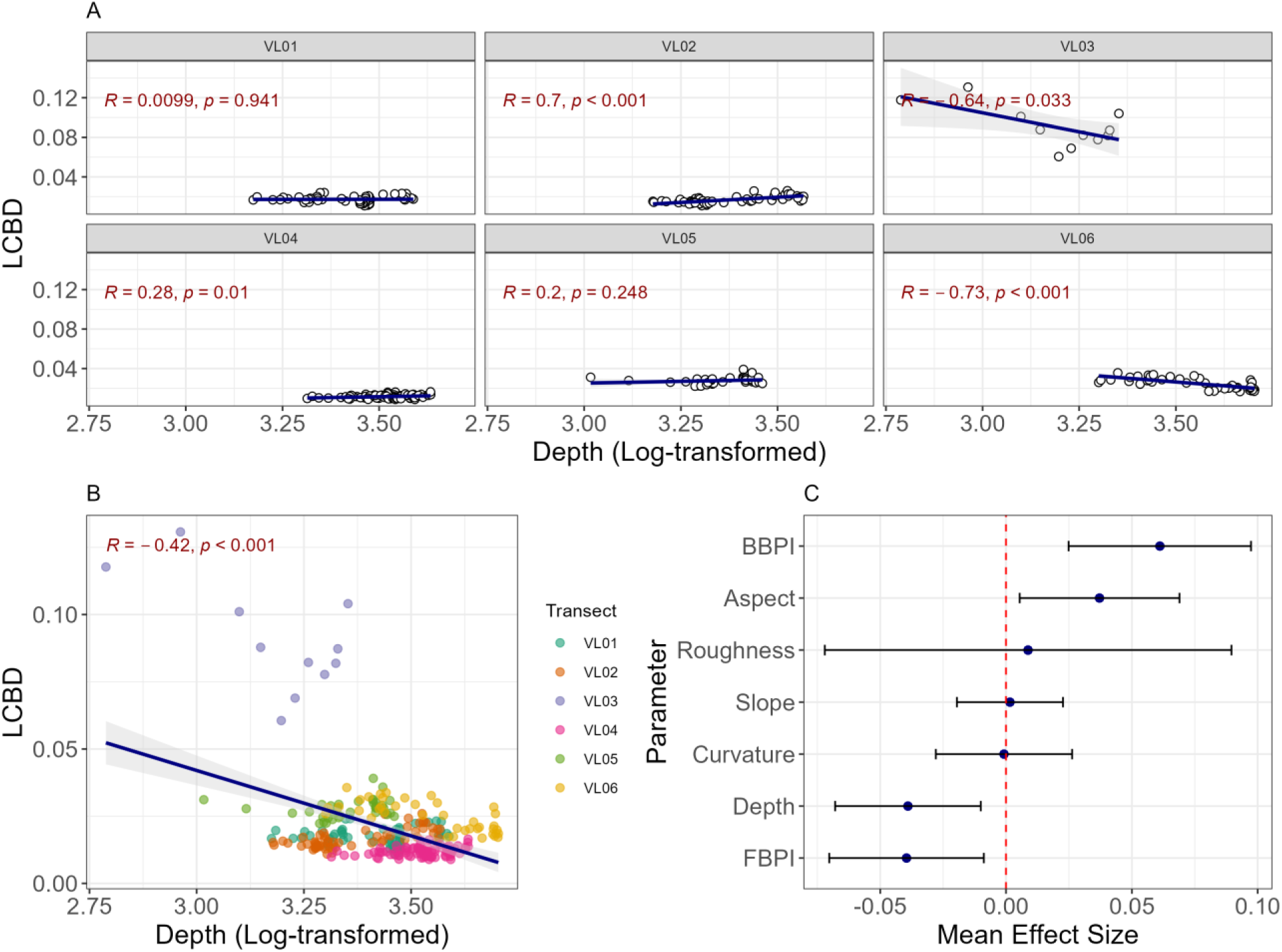
Relationship between local contribution to beta diversity (LCBD) and deep-sea topographic factors. (A) Pearson correlations between LCBD and log-transformed depth within each transect (VL01–VL06; see Table 1). (B) Overall correlation between LCBD and log-transformed depth after pooling data across all transects. (C) Estimated effects of topographic variables on LCBD. Blue dots represent coefficient estimates, and error bars show 95% confidence intervals (CI). Variables with CIs do not cross the red dashed line are considered significant: positive effects to the right and negative effects to the left.

In the mixed effects model, LCBD was negatively associated with depth (*β* = –0.04, *p* < 0.01; 95% CI = [–0.07, –0.01]) and FBPI (*β* = –0.04, *p* = 0.011; 95% CI = [–0.07, –0.01]), while positively associated with BBPI (*β* = 0.06, *p* < 0.001; 95% CI = [0.03, 0.10]) and north-facing aspect (*β* = 0.04, *p* = 0.021; 95% CI = [0.01, 0.07]; Figure 4C).

### Relationships between SCBD and functional traits

Across all transects, SCBD increased significantly with depth-range extent, indicating that species spanning broader depth gradients contributed disproportionately to β-diversity (Figure 5A). Sessility showed a marginally positive association with SCBD (*β* = 0.64, *p* = 0.056; 95% CI = [–0.02, 1.29]), whereas suspension feeders (*β* = –1.10, *p* = 0.012; 95% CI = [–1.96, –0.24]), predators (*β* = –1.16, *p* < 0.01; 95% CI = [–1.98, –0.35]) and filter feeders (*β* = –1.46, *p* < 0.01; 95% CI = [–2.37, –0.54]) were negatively associated with SCBD (Figure 5B).

**Figure 5.**
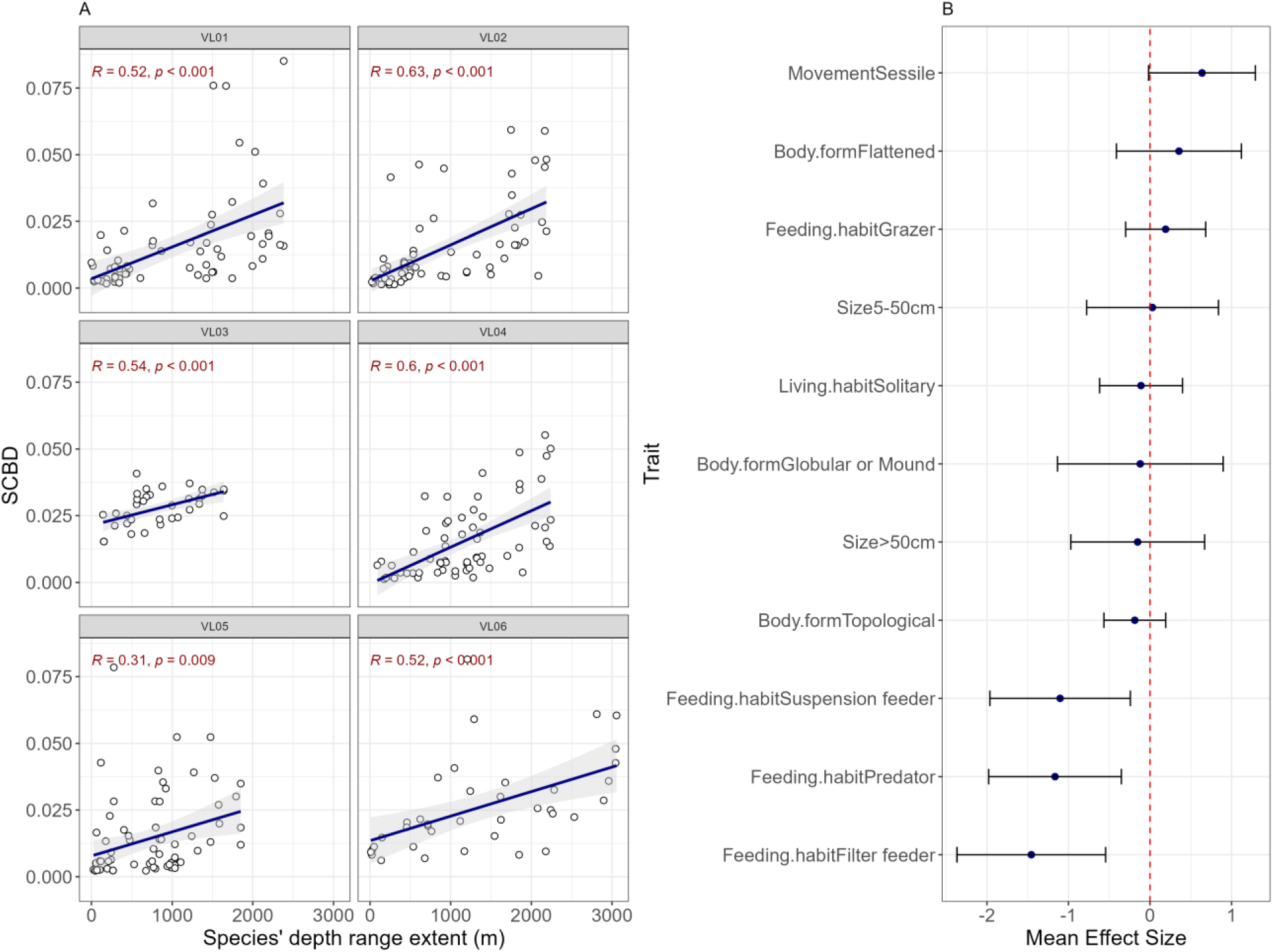
Drivers of species’ contribution to beta diversity (SCBD). (A) Pearson correlations between SCBD and depth-range extent within each transect (VL01–VL06; see Table 1). (B) Estimated effects of life-history and morphological traits on SCBD. Blue dots represent coefficient estimates, and error bars show 95% confidence intervals (CI). Variables with CIs do not cross the red dashed line are considered significant: positive effects to the right and negative effects to the left.

## Discussion

The accelerating expansion of deep-sea resource extraction has brought exceptional attention to seamounts as priority systems for biodiversity assessment and conservation, owing to their structural complexity, high densities of habitat-forming species, and central role in ocean ecosystem functioning (Schlacher *et al*., 2014; Washburn *et al*., 2023). Despite their importance, knowledge of seamount biota remains limited across oceans worldwide, constraining our efforts to effectively protect these vulnerable marine environments (Watling & Auster, 2017, 2021). Our study along the Kyushu–Palau Ridge (KPR) in the western Pacific helps address this gap by integrating species depth-range overlap analysis with β-diversity partitioning to explore how megabenthic community structure varies across depth gradients. We found substantial variation in community structure both within and among seamounts, with nested patterns dominating transects on the western slopes and mixed signatures of nestedness and turnover characterizing the transects on the eastern slopes. Sessile and mobile taxa also differed in their structural patterns, with sessile species typically following nestedness and mobile species showing a mixture of gradual turnover and nestedness. The uniqueness of site-level community composition overall declined with increasing depth and fine-scale bathymetric position index, but increased on north-facing slopes and with broad-scale bathymetric position. Species with wider depth ranges, particularly sessile ones, tended to vary more strongly across depth zones and therefore contributed most to β-diversity. These results identify slope-specific variations, rather than depth alone, as a central feature of KPR megabenthic community organization.

The patterns of depth-range overlap provide important ecological insights into these differences. On the steeper western slopes (Lu *et al*., 2024), assemblages at deeper depths were largely subsets of those occurring in species-rich, shallower portions of the transects, indicating a predominantly nested pattern and relatively ordered community change along the depth gradient. On gentler eastern slopes, the mixture of nestedness and gradual turnover indicates that species were both lost from and replaced along the depth gradient. One possible explanation is that the western slopes exhibit sharper changes in sediment accumulation, substrate type, or local hydrodynamic conditions, whereas habitat conditions may vary more gradually along the eastern slopes (McClain & Lundsten, 2015). These differences are consistent with environmental sorting and habitat-associated species replacement (Leibold & Mikkelson, 2002; Baselga, 2010). Most importantly, distinct community structure persisted between slopes within the same seamount, highlighting the environmental heterogeneity of the seamount ecosystem and cautioning against the application of one-size-fits-all conservation approaches.

The LCBD results reinforce this distinction between broad-scale depth patterns and local habitat variation. We observed an overall negative effect of depth on local contributions to β-diversity when all transects were pooled, which suggests that shallower sites contributed more strongly to the overall compositional variations across the region. This pattern is broadly consistent with productivity theory: declining particulate-organic-carbon flux with depth can reduce the energy available to benthic communities and lead to more similar assemblages at depth (Clark *et al*., 2010). However, the relationship was not consistent across slopes, indicating a clear slope dependency. Thus, the observed negative relationship at the regional scale should be interpreted as a general trend rather than a deterministic effect of depth. The positive relationships on VL02 and VL04 are particularly noteworthy because they show that local habitat conditions can override the expected productivity-related pattern. Cordes et al. (2025) similarly found that deep-sea chemosynthetic and cold-water coral communities did not consistently exhibit declining abundance or diversity with depth. Instead, localized energy sources, nutrient retention, species adaptations, and geological or biogenic habitat heterogeneity helped maintain high biodiversity across depth ranges. In our study, the contrasting LCBD patterns may likewise reflect differences in the relative contribution of slope-specific hydrodynamics and habitat heterogeneity (Tracey *et al*., 2011; Bishop *et al*., 2022; Shen *et al*., 2024). Aspect provided an additional indication that local physical setting influences community composition. North-facing sites had higher LCBD, consistent with regional observations that deep currents passing through the canyons separating KPR seamounts intensify bottom flows along the corresponding north-facing slopes (Wang et al., 2023; Zhou et al., 2022). Enhanced flow could increase the delivery of particulate organic matter to suspension-feeding organisms and influence the transport of larvae, which may result in the greater compositional differentiation of these sites. Although we were unable to measure these factors directly, they provide a plausible explanation for such transect-level contrasts.

These results have direct implications for conservation planning. Shallow habitats on the steeper western slopes are important conservation targets because they may contain unique assemblages and contribute strongly to observed compositional differentiation. Similar patterns could be found in the Eastern Pacific, where communities on slopes steeper than 30° showed greater compositional similarity and lower β-diversity (McClain & Lundsten, 2015). On the gentler eastern slopes, the presence of deeper compositionally distinct assemblages indicates that an effective conservation strategy should encompass multiple depth zones.

Species-level contributions to β-diversity provided a complementary perspective. Wide-ranging generalists contributed more strongly to β-diversity than narrow-ranging specialists or rare species, reflecting their greater variation across sites. It does not, however, mean that these taxa should automatically be treated as conservation priorities. High SCBD can reflect abundance and broad occupancy, whereas rare or endemic species may have low SCBD despite being highly vulnerable (Legendre & De Cáceres, 2013; Rodríguez-Lozano *et al*., 2023). High-SCBD taxa are therefore useful indicators of broad-scale community change, while conservation decisions should additionally consider rarity, endemism, habitat vulnerability, functional importance, and taxonomic uncertainty.

To sum up, our findings reveal that KPR seamount megabenthic communities are shaped by the interaction of broad bathymetric gradients with slope-specific habitat structure. Western slopes were characterized primarily by nestedness, whereas eastern slopes contained stronger evidence of species replacement along the depth gradient. These findings emphasize the importance of considering depth and slope-specific variation when assessing the conservation of deep-sea seamount communities.

## Data availability statement

The data that support the findings of this study are available in Figshare at https://doi.org/10.6084/m9.figshare.31320511.

