## supplementary material for "Contrasting megabenthic community structure across western Pacific seamounts to ∼5,000 m depth: Implications for conservation"

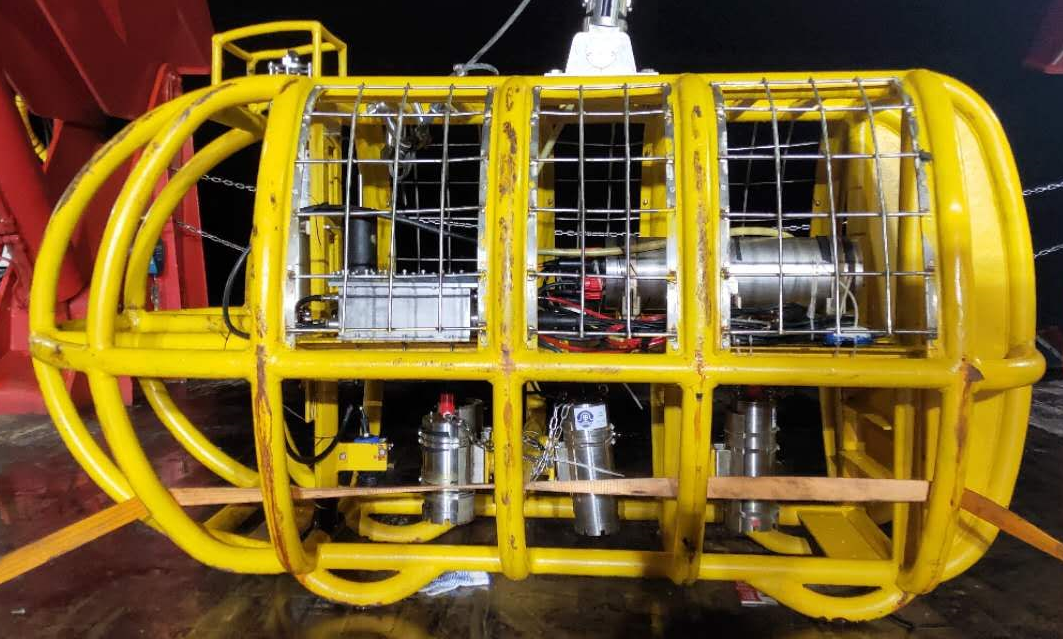

Figure S1. Survey equipment: The towed camara system used for imaging surveys, capable of operating at depths up to 6,500 m.

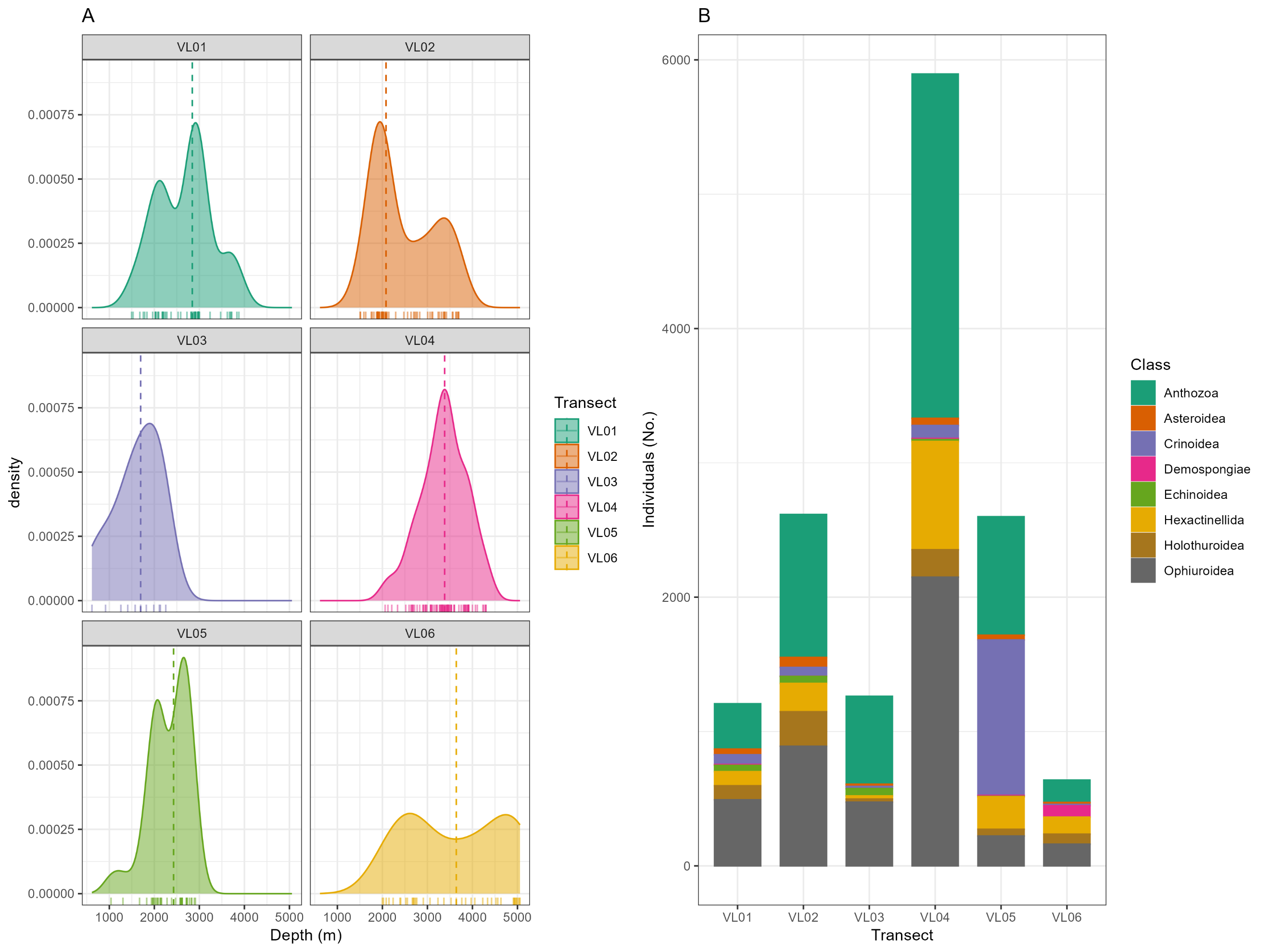

Figure S2. Depth (A) and species (B) profiles of each surveying transect. In panel (A), marginal rugs along the x-axis represent the precise depths of individual sampling plots.

Table S1.

The observed and expected overlap patterns within each transect as per community type. Transects on the same seamount are displayed with the same color.

| Transect | Slope | Community type | Observed overlap triplet | Expected overlap triplet | Signs |
| --- | --- | --- | --- | --- | --- |
| VL01 | Western | Full | (442, 463, 1111) | (672, 672, 672) | $\left( -,-,+ \right)$ |
|  |  | Sessile | (173, 151, 456) | (260, 260, 260) | $\left( -,-,+ \right)$ |
|  |  | Mobile | (55, 88, 133) | (92, 92, 92) | $\left( -,-,+ \right)$ |
| VL02 | Eastern | Full | (119, 884, 1412) | (805, 805, 805) | $\left( -,+,+ \right)$ |
|  |  | Sessile | (53, 364, 573) | (330, 330, 330) | $\left( -,+,+ \right)$ |
|  |  | Mobile | (13, 112, 175) | (100, 100, 100) | $\left( -,+,+ \right)$ |
| VL03 | Western | Full | (62, 190, 378) | (210, 210, 210) | $\left( -,-,+ \right)$ |
|  |  | Sessile | (25, 76, 152) | (84.3, 84.3, 84.3) | $\left( -,-,+ \right)$ |
|  |  | Mobile | (7, 15, 56) | (26, 26, 26) | $\left( -,-,+ \right)$ |
| VL04 | Eastern | Full | (270, 761, 1049) | (693, 693, 693) | $\left( -,+,+ \right)$ |
|  |  | Sessile | (103, 175, 388) | (222, 222, 222) | $\left( -,-,+ \right)$ |
|  |  | Mobile | (22, 166, 190) | (126, 126, 126) | $\left( -,+,+ \right)$ |
| VL05 | Eastern | Full | (444, 739, 1232) | (805, 805, 805) | $\left( -,-,+ \right)$ |
|  |  | Sessile | (175, 272, 499) | (315, 315, 315) | $\left( -,-,+ \right)$ |
|  |  | Mobile | (28, 134, 163) | (108, 108, 108) | $\left( -,+,+ \right)$ |
| VL06 | Western | Full | (175, 207, 321) | (234, 234, 234) | $\left( -,-,+ \right)$ |
|  |  | Sessile | (76, 68, 132) | (92, 92, 92) | $\left( -,-,+ \right)$ |
|  |  | Mobile | (13, 35, 43) | (30.3, 30.3, 30.3) | $\left( -,+,+ \right)$ |
